# *In Silico*, Brain Mesh Platform for Computing Topographic Dependent Internal Facets

**DOI:** 10.1101/2022.10.27.514138

**Authors:** Caleb Shaw, Cecile Riviere-Cazaux, Dylan Obrochta, Stephanie Otto, Lawrence Ray, Lars Strother, Terry Burns, M. Rashed Khan

## Abstract

Individualized and anatomically correct computational models of the brain can be leveraged to improve knowledge of drug dispersal following simulation of drug delivery. Using a patient’s magnetic resonance image (MRI) scans, we were able to reconstruct the pial surface of the brain of the left hemisphere with strong anatomic accuracy. We then established the major internal features, including the lateral ventricle, a tumor, and drug delivery catheters. These were able to include relevant tissue characteristics such as porosity and permeability in the Multiphysics platform COMSOL to create a platform for brain modeling. To test the performance of this platform, we simulated direct drug infusion in both a healthy patient brain and a diseased patient model, focusing on glioblastoma (GBM). Using this platform, we simulated perturbed convection enhanced delivery of a cancer medication (similar to temozolomide (TMZ) but modeled using methylene blue) to the tumor. Consequently, with our patient derived model, we are able to simulate solute dispersal and fluid flow representative of *in vivo* conditions.

## Introduction

Glioblastoma is the most common primary brain tumor that is currently incurable and inevitably fatal [1]. Despite maximally safe resection and aggressive chemoradiation, the prognosis for patients with this disease remains dismal at a mere 12 to 18 months [2]. No drug has improved survival for patients since temozolomide in 2005. While certain drugs, such as carboplatin and paclitaxel, have shown potency in preclinical models, these effects have failed to translate clinically. Such difficulties are due in part to the limitations of delivering drugs past the blood-brain barrier and achieving an effective intratumoral dose without off target deleterious sequelae on adjacent normal tissue [3]. Convection enhanced delivery (CED) is a method of direct intratumoral drug injection to bypass the blood brain barrier (BBB), resulting in substantially higher dosing within the tumor when compared to systemic drug administration [4]. However, to date, CED has not substantially improved patient outcomes [5]–[7], with prior work suggesting that this lack of success may be due to non-optimal CED techniques, including suboptimal catheter placement, catheter inner and outer diameters, flowrates, and flow scheduling [5].

Optimization of CED treatment across patients has been difficult due to the brain’s complex topography and its variability across patients, in addition to heterogeneous tumor shapes and tissue characteristics. A promising potential solution to improve drug distribution via CED is treatment simulations via *in-silico* individualized modeling from each patient’s pre-operative MRI prior to catheter implantation. In this manner, treatment plans, including catheter location, flow rate, and duration of drug delivery, could be devised on a patient-to-patient basis to maximize the tumor volume covered by the drug. Consequently, multiple efforts have been made to computationally simulate CED of candidate therapies into tumor and brain [8], [9]. However, a limitation of these simulations is that the brain topographies are unable to accurately replicate that of the outermost layer of the cerebral cortex that can be approximated by reconstruction of the pial surface. Many factors impact solute dispersal within the brain including interstitial fluid flow (IFF), local tissue variations including the capillary action of white matter tracts, and local temperature variations. These parameters occur within and are impacted by the spatial arena in which it occurs—being the pia mater. Consequently, constructing a patient’s parenchymal topographic structure is a necessary step in creating a replicating interstitial brain simulation that can be deployed in diffusion studies to determine the impact of each individual’s response with molecular delivery, IFF, pressure, and tumoral micro-structure induced perturbations.

Established in our prior study, by Shaw et al. (“Convection-enhanced diffusion and directed withdrawal of methylene blue in agarose hydrogel using finite element analyses”), we have shown that CED-mediated molecular dispersion can be improved by utilizing an additional extraction/outflow catheter to induce a fluid “conveyor belt” from the inflow delivery catheter to the extraction/outflow catheter that acts as a “sink” by locally depleting solutes. Within brain tumor simulation models, drug delivery will additionally be impacted by anatomic perturbations, such as sulci and gyri, in addition to increased and heterogeneous intra-tumoral density. We hypothesize that simulations including a highly accurate pial surface topography, particle diffusion, and fluid convection can be utilized to test and optimize CED conformations with more accuracy than previously afforded by other brain models.

As such, given the promise of CED for drug delivery despite its limited success to date, there exists an unmet need to optimize CED through simulations through incorporation an individual’s brain topography and inflow/outflow catheter placement. To address this gap, we present an *in silico* individualized brain model with a topographically-reconstructed pial surface from a patient’s MRI and evaluate its performance using a CED simulation (Figure 1). After converting the MRI obtained from the Dallas Lifespan Brain Study repository and converting it to a CAD object per our prior study by Ray et al. (“Synthetic Brain Fabricated from MRI, Using Agarose-gel for Proof of Concept Diffusion Studies on Tissue-Substitutes”), a mesh reconstruction of the countersurface of the image was established using a volumetric based neuro-reconstructing software-FreeSurfer -- to create the FreeSurfer mesh (Figure 2). After reconstruction, the mesh was edited in Meshmixer and inputted into COMSOL Multiphysics. An *in-silico* platform was then created utilizing individualized topographic data and parameterized with literature-derived tissue data, with our pre-established CED simulation inputted into COMSOL. In our disease model, a GBM was created and applied to the model’s base platform in a spherical shape. In conclusion, with our normal and diseased brain models, we present an improved and topographically accurate, MRI-derived brain mesh model for CED simulation and drug diffusion studies.

**Figure 1:**
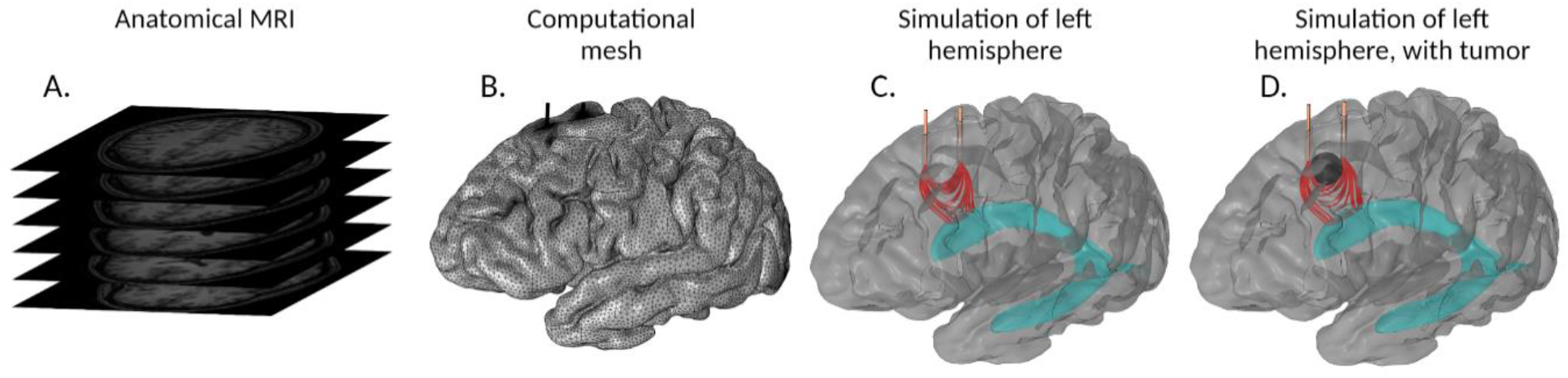
Workflow for establishment of the *in-silico* brain mesh platform. A. MRI stack of axial plane slices which are utilized to build the computational mesh. B. COMSOL Multiphysics computational mesh of a left hemisphere reconstructed from the anatomical MRI in 1A. Two catheters were implanted in healthy brain to simulate local drug delivery. C. COMSOL Multiphysics velocity simulation of catheter injecting solute into healthy brain. The left catheter performed infusion while the right catheter extracted at the same rate D. COMSOL Multiphysics velocity simulation of local drug delivery via implanted catheters.

**Figure 2:**
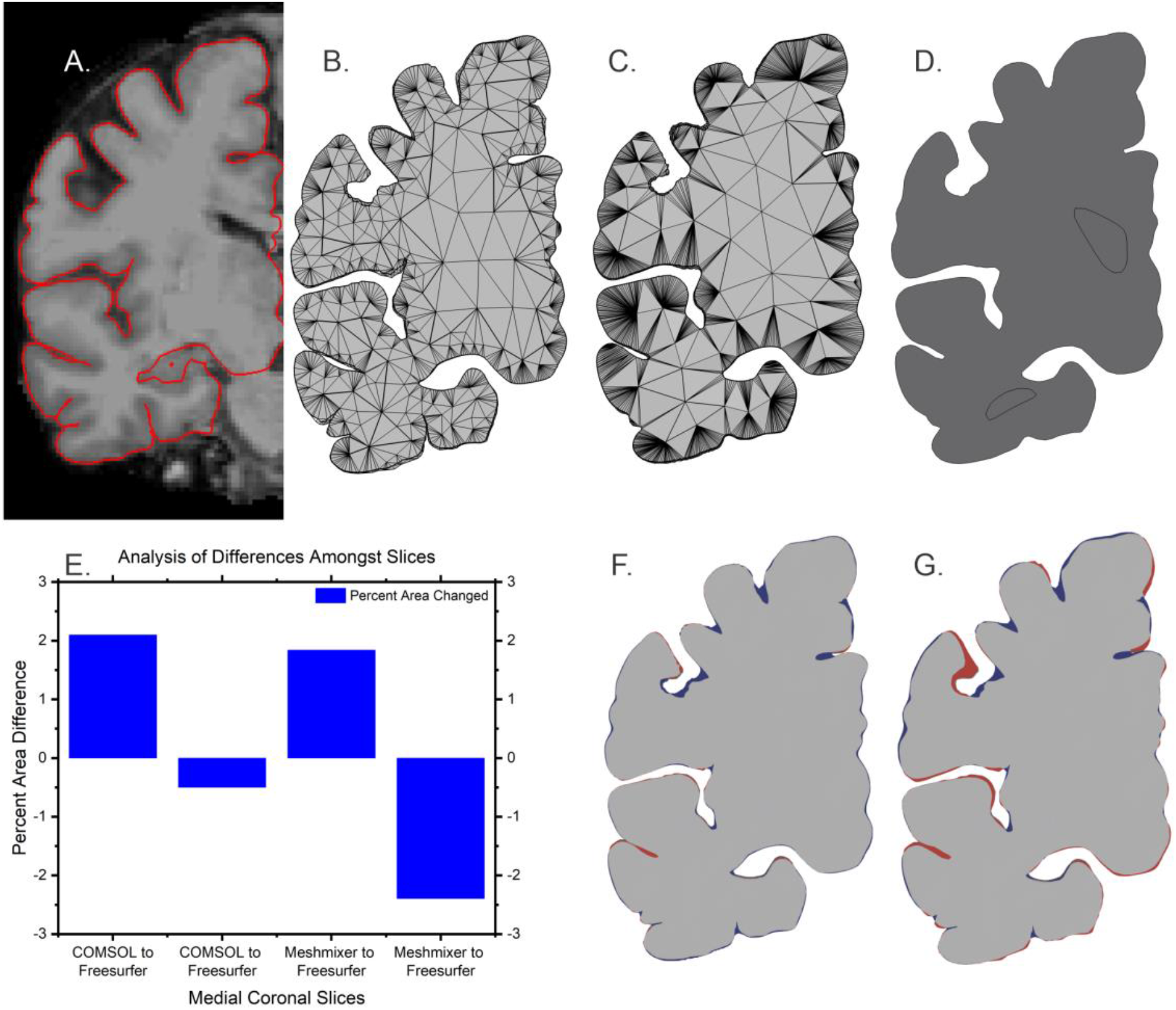
Topographic Replication using an individual’s MRI data. Coronal slice, from the middle of the brain, comparison of pial surface of meshes to demonstrate conformity of replication during processing. A. MRI slice with pial surface highlighted in red. B. FreeSurfer slice generated by FreeSurfer. C. The mesh slice after the edits made in Meshmixer. D. COMSOL Multiphysics computational mesh after simulation. E. Plot of area added and subtracted to the original FreeSurfer slice. F. Slice showing area added and subtracted to original FreeSurfer mesh by the Meshmixer edited mesh. The blue regions show areas added and the red show areas subtracted from the FreeSurfer slice. G. Slice showing the blue added and the red subtracted regions of the FreeSurfer slice comparing to the COMSOL slice.

## Methods and Materials

### MRI data

Open-source MRI data was acquired from the NeuroImaging Tools and Resources Collaboratory, the Image Repository from the Dallas Lifespan Brain Study http://www.nitric.org/. A T1 weighted scan was utilized from a 73-year-old female subject, who was imaged on a 3T Siemens ECAT HR MRI scanner utilizing MPRAGE (magnetization-prepared rapid gradient-echo).

### MRI to Volumetric Data

The neuro-analytic/reconstruction software FreeSurfer was utilized to reconstruct the nifty (neuroimaging informatics technology initiative) file and to establish a mesh render of the brain [10]. The mesh output was converted to an STL file type with the software file converting command.

### Modification of Volumetric Mesh to Usable Mesh

MRIs utilize magnets to induce polarization of the magnetic poles in hydrogen atoms in water. The decay of the polarized orientation of the hydrogen allows the MRI to interpret regions of high and low water content inside tissue. FreeSurfer interprets this MRI data to develop a mesh replicating the brain’s volume in an interpretative process recognized by industry. This operates by comparing the new scan to a standard brain which has its brain regions defined. Based on relative voxel intensities, FreeSurfer identifies certain white matter locations and extrapolates from these points the sections of the brain. Our objective was to integrate the FreeSurfer generated mesh into COMSOL for interstitial solute computation. Importing the file generated by FreeSurfer into COMSOL in the stereolithography (stl) format generated numerous geometry and mesh creation errors. These errors were generated by the non-manifold construction of the original object created by FreeSurfer, in that the surface was incomplete and did not fully encapsulate an interior volume. We manipulated the surface in Meshmixer by Autodesk to identify the errors and to make the surface manifold (complete as a solid objected containing a volume) [11]. Meshmixer hosts tools such as adjustments to the number of meshing elements, total object smoothing features, brushes to adjust specific sites, and remeshing to distribute mesh elements equally throughout the object. Once manifold, we successfully imported the object into COMSOL from Meshmixer to identify areas in the mesh needing further refinement as indicated by COMSOL. The object was reimported into Meshmixer to adapt it with the available tools utilizing feedback from COMSOL, including smoothing brushes and volume adjustments (increasing floors of sulci and lowering peaks of gyri). This process between COMSOL and Meshmixer was iterated until COMSOL generated a complete mesh without construction errors. Our iterative process required approximately 100 trials to reach successful computational meshing. This final mesh generation was calibrated for fluid dynamics adjusting the meshing size to be appropriate for fluid flow.

### COMSOL Simulation

Once the brain MRI was constructed into a usable mesh, we imported it into COMSOL Multiphysics [12]. COMSOL Multiphysics then re-meshed this usable mesh into a computational mesh that can resolve differential equations. The “normal” meshing setting was utilized when defining the brain tissue in the meshing of the Tumor and No Tumor models, while the catheters and associated structures were meshed as “Extra Fine.” The associated structures were cylinders that formed a sleeve around the catheters to aid COMSOL’s computational meshing. The sleeves acted as steps for meshing sizes from the very small elements within and surrounding the catheters to the large elements within the brain’s courser further afield features. COMSOL Boolean combination function sliced the sleeves’ top so that the face of the sleeves matched the topography of the brain. Furthermore, the sleeve interface with the rest of the brain was not defined as a boundary but a continuous brain matrix. The change induced is that of more mesh elements with more data points calculated within a unit volume of the sleeves than in the surrounding brain. An additional volumetric shape corresponding to the ventricular system was added to provide major internal features to the model [13]. The physics used within COMSOL were the Transport of Diluted Species to simulate mass transport and the Free and Porous Media Flow module for momentum fluid flow. Specific properties used in the simulation are shown in Table 1.

**Table 1.** Properties Used in COMSOL Multiphysics

| Property | Value | Units |
| --- | --- | --- |
| Particle Radius (Methylene Blue) | $7 \times 10^{-10}$ | [m] |
| Input and Output Flowrates | $3 \times 10^{-6}$ | [L/min] |
| Tortuosity of 0.6% agarose/brain | 1.6 | [14] |
| Catheter Inner Radius | 0.43 | [mm] |
| Catheter Outer Radius | 0.73 | [mm] |
| Catheter Height | 22 | [mm] |
| Porosity of 2% agarose | 0.831 |  |
| Porosity of 0.6% agarose | 0.951 |  |
| Tumor Biomarker Initial Concentration | $3*10^{-9}$ | [mol/m<sup>3</sup>] |
| Drug Input Concentration | $3*10^{-1}$ | [mol/m<sup>3</sup>] |
| Tumor Biomarker Rate of Generation | $1*10^{-7}$ | [mol/(s*m<sup>3</sup>)] |
| Catheter Separation | 16 | [mm] |
| Tumor Diameter | 13 | [mm] |
| Water Viscosity | $1.002*10^{-3}$ | [kg/(m*s)] |
| Boltzmann's Constant | $1.38*10^{-23}$ | [((m<sup>2</sup>)*kg)/(K*s<sup>2</sup>)] [15] |
| 0.6% Agarose Pore Size | 794.18 | nm<sup>2</sup> |
| 2% Agarose Pore Size | 381.754 | nm<sup>2</sup> |
| Temperature for material properties | 293.15 | K |

COMSOL Multiphysics takes differential equations and arrives at an approximate solution through finite element methods (FEM). Three sets of differential equations are utilized: 1) the Navier-Stokes Equations used for laminar flow through the catheters [16], 2) the Brinkman Porous Flow Equations for flow through the porous hydrogel [17], and 3) Fick’s Second Law of Diffusion for diffusion of the solute, listed as Equation 1, Equation 2, and Equation 3 respectively [16], [18]. Equation 4 defines the Convection Diffusion equation utilized by COMSOL Multiphysics to combine fluid flow and diffusion [19]. Diffusivity and permeability were the two physical properties that defined solute motion within the material, defined as the Stokes Einstein Diffusivity and the Kozeny-Carman Permeability (Equations 5 and 6, respectively) [15], [20]. Within the equations, the terms include:

Boltzmann constant (k_b_), Brinkman flow (Q_Br_), concentration (c) or (c_i_), Darcian permeability (k_1_), diffusion flux (J) or (J_i_), coefficient of diffusion (D) or (D_i_), fluid density (ρ), fluid dynamic viscosity (μ), fluid pressure (p), fluid velocity (u), external applied forces (F), forchheimer drag coefficient (β_F_), gradient vector (▽), identity vector (I), mass source or sink of species i, particle radius (R) or (R_i_), pore diameter (d_pore_), porosity (ε) and (α), pressure (p), stress tensor (T_s_), temperature (T), and time (t) [15]–[17], [21].

#### Navier-Stokes Equation

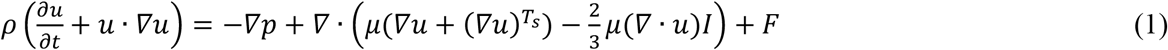

#### Brinkman Porous Flow Equation

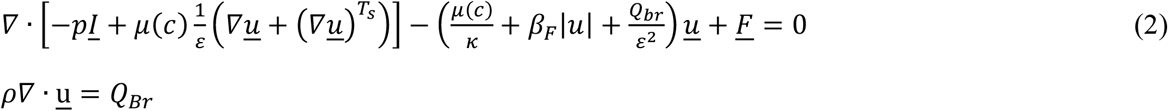

#### Fick’s Second Law of Diffusion

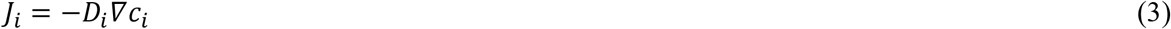

#### Convection Diffusion Equation

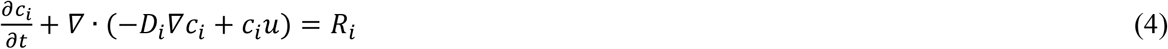

#### Stokes Einstein Diffusivity

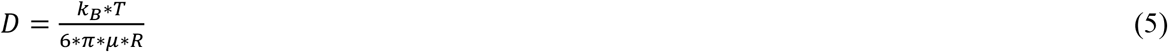

#### Kozeny-Carman permeability

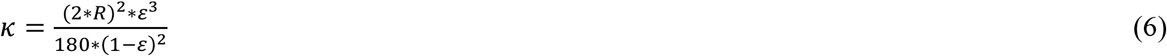

#### Porosity Permeability Pore Size

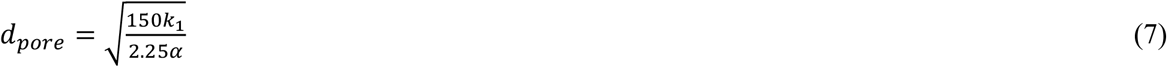

### Brain Tissue Properties Approximated by Agarose Hydrogels

Previous efforts have suggested that brain tissue diffusivity properties are most accurately replicated in 0.6% agarose hydrogel [22], [23]. With agarose hydrogels, prior studies have established the permeability, pore size, and the relation of these two properties to porosity, defined in Equation 7 [24]–[26]. Furthermore, 2% agarose can approximate dense regions of GBMs [27], [28].

## Results

With the Meshmixer’s brushing and meshing number tool, the computational mesh data errors were individually corrected. In COMSOL Multiphysics, the finite elements use mesh elements to resolve the differential equations requiring a continual element to element sequence covering the entire computed geometry. Consequently, the computational meshing in the COMSOL software, was sensitive to construction and fails in high aspect conformations. Upon successful computational meshing, the Tumor and No Tumor Model created 10^6^ meshing elements as seen in Supplemental Table 1.

Importation of the coronal slices from the original MRI into FreeSurfer resulted in creation of FreeSurfer volumetric data (Figure 2A-B). We compared edited files to the FreeSurfer file rather than the direct MRI since the FreeSurfer file defines the pia mater from the MRI in a usable format. The FreeSurfer slice was then iteratively edited between Meshmixer and COMSOL, resulting in a final edited Meshmixer slice (Figure 2C) which could be inputted into COMSOL as a post-computational mesh after simulation (Figure 2D). The coronal slices were taken from the medial section of the brain while the anterior and posterior slices are shown in supplemental (Supplemental Figure 1). From visual inspection, the sharp interior curves and features of the FreeSurfer mesh slice (Figure 2B) had been smoothed in the Meshmixer and COMSOL mesh slices. From the anterior, medial, and posterior slices, the slice with the greatest change between the FreeSurfer mesh slice to the final COMSOL mesh was the most posterior section (Supplemental Figure 1I-K) with the semi-disjointed portion in the top left being fully merged into the main body in the Meshmixer edited and COMSOL surface resulting in 2.5% difference in area. Despite these blurred details, this computational mesh construction closely replicates the original MRI volumetric data.

To quantify the impact of processing between Freesurfer, Meshmixer, and COMSOL on the volume of the slice, we subtracted the Freesurfer medial coronal slice from the edited Meshmixer slice (Figure 2B-C) in Autodesk Fusion360 [29]. This subtraction gave a sweep of the intermediate areas differentiating the slices. This was process was repeated between the FreeSurfer slice and the COMSOL slice (Figure 2B and D). Areas added and subtracted to the FreeSurfer slice in order to achieve the Meshmixer (Figure 2F) and COMSOL (Figure 2G) slices were depicted in blue and red, respectively. The total area changes from the baseline FreeSurfer mesh were utilized to compute the area differential (percent change) as compared to Meshmixer and COMSOL (Figure 2E) for a slice. Furthermore, we performed this process with the volumes of the total meshes to determine the total volume change. The Meshmixer edited mesh volume was 3.1% different from the FreeSurfer mesh and the COMSOL surface mesh volume was 4.1% different from the FreeSurfer mesh, illustrating the conserved volume of our meshes throughout processing.

### Convection Enhanced Delivery Simulation

To test our *in-silico* brain mesh model, we performed a simulation of convection enhanced delivery into the brain, both healthy and with a tumor in the disease model. Unlike conventional CED models, an extraction/outflow catheter was included to hypothetically perturb fluid flow and direct solute dispersion. With a diffusion only process, we would expect concentration profiles to follow an error function decline of Fickian diffusion. Two simulations were performed: 1) drug delivery into a normal brain and 2) drug delivery into a disease model (GBM), (Figure 3A and B). Line scan diffusion plots were used to quantify the drug diffusion profile along a sagittal cut of the brain 2 mm below the catheters and above the tumor center. The catheters were placed at 6 mm for infusion and at 22 mm for extraction (Figure 3F). Furthermore, in Figure 3C and 3D, the maximum intensity of the profile was reduced to one-third of the input concentration to allow visualization of the drug’s interaction with the tumor mass. Drug delivery in the healthy brain model revealed two disruptions in molecular diffusion, one induced by the extraction catheter and the other by the sulci occurring near the 0 mm mark on the plot. In our model, extraction/outflow catheter constructs a “conveyor belt” of flow from the injection catheter to the extraction catheter. This set-up induces preferential drug distribution to favor the direction of the extraction catheter that acts as a solute “sink.” Additionally, the presence of sulci induced the drug to accumulate around the edge of the pia mater which locally increases the drug concentration, as seen visually in Figure 3A with the drug concentrating around the edges of the sulcus.

**Figure 3:**
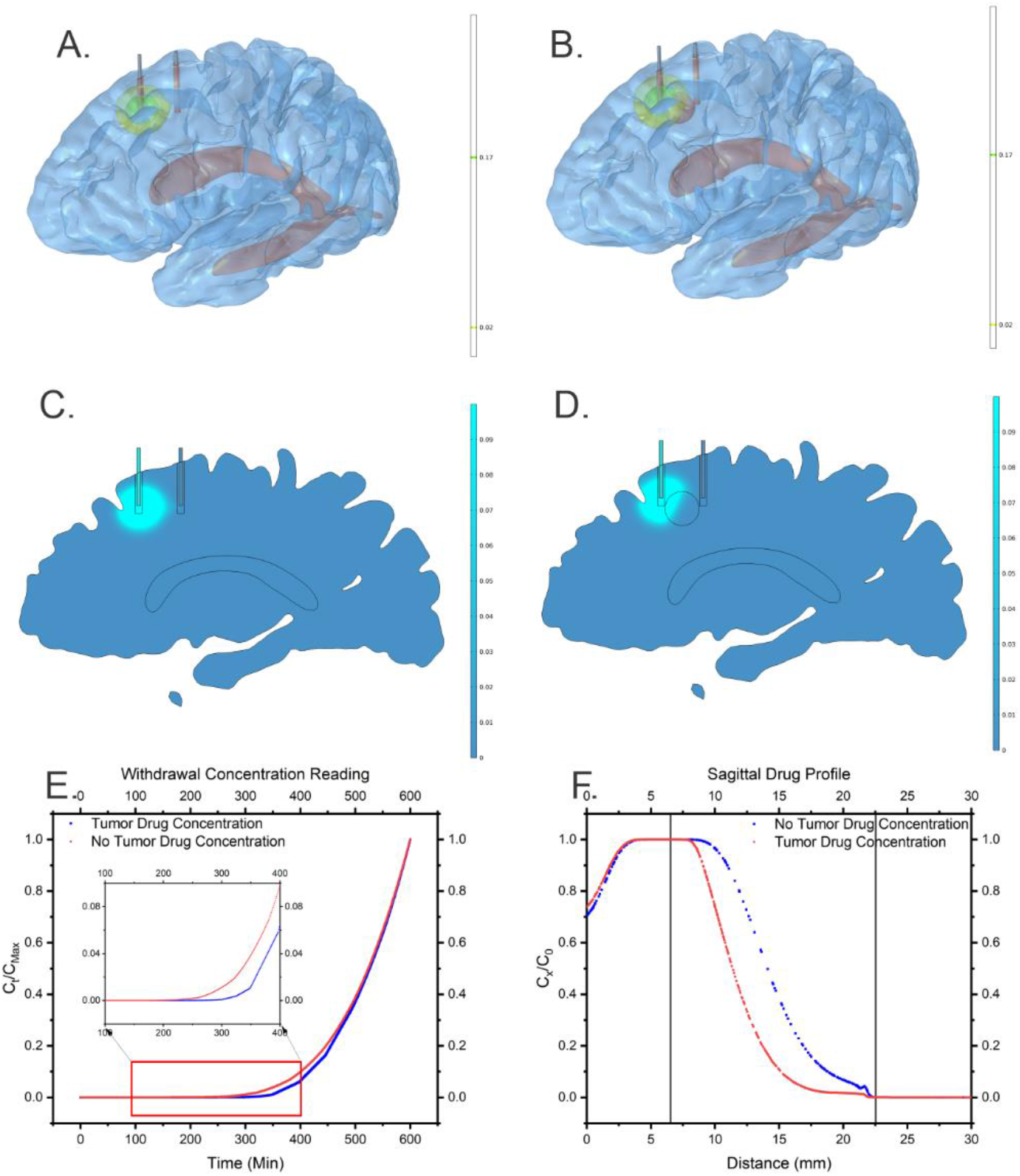
Simulation results of CED: A. Concentration profile of drug delivered from input catheter to extraction catheter in the No Tumor Model.. B. Concentration profile of drug delivered from input catheter to the extraction catheter across the tumor construct in the Tumor Model. C. Sagittal slice showing the concentration profile of drug delivered from input catheter to extraction catheter in the No Tumor Model. The profile is adjusted so the maximum is one-third of initial concentration injected. B. Sagittal slice showing the concentration profile of drug delivered from input catheter to the extraction catheter across the tumor construct in the Tumor Model. The profile is adjusted so the maximum is one-third of initial concentration injected. D. Concentration profile of drug of nutrient present in the ventricle dispersing throughout the brain from the ventricle in the Tumor Model. The profile is adjusted so the maximum is one-third of the initial concentration present in the ventricle. E. Normalized concentration profile of bio-marker on a sagittal line across the entire brain at the catheter tip. F. Normalized concentration profile of delivered drug on a sagittal line across the entire brain at the catheter tip. Blue line is the profile from the No Tumor model and the red line is from the Tumor Model.

We hypothesized that in the disease model, presence of the dense tumor mass would disrupt fluid flow following focal delivery, with the increased intra-tumoral density resulting in preferential fluid flow to the areas around the tumor that have lower tissue density. Indeed, in our simulation, the zone within the tumor mass (approximately 10-20 mm on the chart), the drug concentration dropped steeper relative to the healthy model (Figure 3F). Notably, obstruction of fluid flow by the tumor further disrupted drug distribution near the sulci at the 0 mm mark – as fluid could not flow past the tumor. It instead preferentially distributed around the sulcus, as seen by increased drug concentration at this point compared to the healthy brain model (Figure 3F). From these plots, the recovery of drug abundance in the extraction/outflow catheter over time was determined. When comparing the recovery drug concentration between the healthy and GBM model, the diseased simulation demonstrated decreased drug abundance from 300-500 minutes, likely owing to higher intra-tumoral density limiting fluid flow (Figure 3E).

### Biomarker recovery in a GBM model

In addition to drug delivery, biomarker abundance is of particular interest in GBM simulation models to determine the expected recovery at various distances from the tumor, particularly in phase 0 signal-finding studies (Fig. 4). Toward this goal, we simulated biomarker recovery from the extraction/outflow catheter in our GBM model. CED manipulates the biomarker profile. Biomarker recovery can elucidate concentration levels through the surrounding tissue. Within this model, the tumor core (∼15 mm) is set to have an initial maximal biomarker concentration which then decays at increasing distances from the tumor core, assuming that the most aggressive portion of the tumor with highest tumor cellularity is likely to produce the greatest concentration of tumor-derived biomarkers. We quantified from the extraction/outflow catheter and demonstrated a linear increase over time, potentially due to induced perfusing flow from CED and increasing biomarker concentration from generation relative to baseline. The inflow and outflow disrupted the profile but utilizes the conveyor belt to sample throughout the intermediate tissue relaying additional information in excess of a single point extraction which induces a local sink.

**Figure 4:**
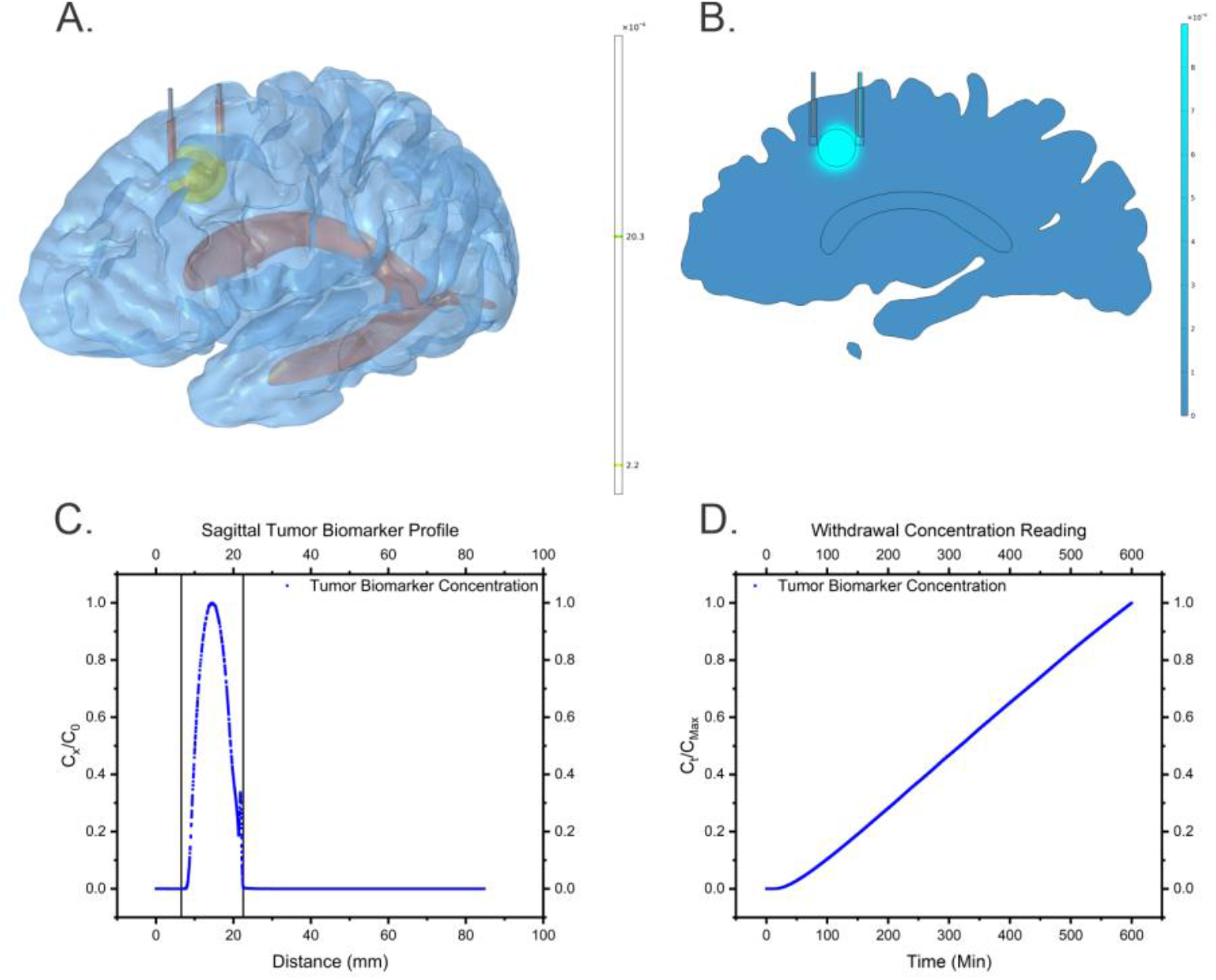
Concentration profiles tumor biomarker. C. Biomarkers generated within the tumor being extracted from extrusion catheter. The tumor mass had an initial concentration present upon the start of the simulation followed by species generation throughout the tumor mass. The profile is adjusted by the maximum shown being the initial bio-marker concentration increased by 3*10^5^.

### Multi-Catheter and Variable Flow Infusion

Having performed the drug delivery and biomarker recovery in a 2 catheter CED system, we then asked the question of how our model would perform when evaluated with more elaborate catheter and flow scheduling. We built our full healthy and GBM models with four catheters surrounding a tumorous mass. As in the CED model, we simulated drug delivery in the healthy brain (Figure 5A) and GBM model (Figure 5B) for 10 hours.

**Figure 5.**
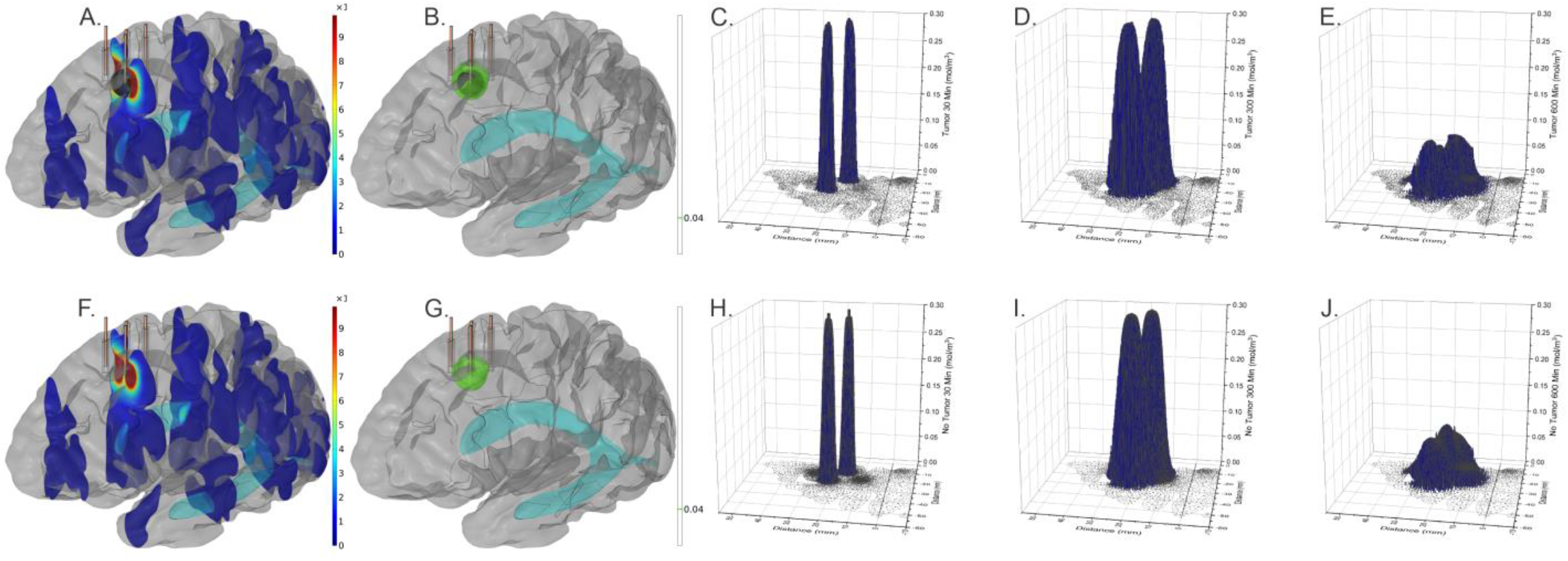
Four Catheter Variable Flow Simulation. A. Diseased simulation with a velocity magnitude shown through coronal slices illustrating induced flow from the catheters. B. Diseased model simulation with yellow and green drug concentration iso-curves. The model is the same as the healthy model except for inclusion of tumor mass seen as the black sphere between the two catheters. C., D., and E. show the normalized concentration profile of input drug at 60-minute, 300-minute, and 600-minute time stamps in the GBM model. F. Healthy model simulation showing induced velocity profile from catheter flow. G. Healthy model simulation with green and yellow iso-curves marking the concentration of drug delivered. Yellow iso-curve surrounds the input catheter on the left-side and shows perturbation to the output catheter on the right side. Lower turquoise surface volume is the ventricle. H., I., and J. show normalized drug concentration profile at 60-minute, 300-minute, and 600-minute time stamps in the healthy model. The x-y plane was taken half-way between the catheter tips and the center of the tumor mass (2 mm below catheter tips and 2 mm above tumor center of mass). Notable on figure J is the leveled concentration profile over the tumorous region.

We constructed a 4-catheter system to evaluate optimal drug delivery. The catheters “box in” the drug to maximize dosing in the tumor and limit dosing in healthy tissue. For this situation, we did not include a central catheter so as to emphasize a boxing effect from the surrounding catheters. Instead we offset the catheters so 2 are closer to the tumor center while the other 2 are further out. With this configuration, we launched a variable flow scheme. The first 300 minutes, the inner catheters infused at a constant rate while the outer catheters withdrew at the same rate. With this scheme, the drug would undergo a conveyor belt to spread through the tumor. For the second 300 minutes, the inner catheters conducted withdrawal while the outer conducted water infusion without drug. The objective being the inner catheters removing excessive drug dosing on the side away from the tumor while the outer catheters would push tumor side drug back through the tumor. The concentration profile of the drug in the normal and GBM model was evaluated on an axial plane 2 mm below the catheter tips at 30, 300, and 600 minutes into drug delivery (Figure 5 C-E and H-J). The 30-minute profile shows the sharp decrease soon after starting infusion being relatively similar. The decreased tumor penetrability is notable in comparing Figure 5 D and I with the dip in concentration between the two infusion points. Meanwhile, after the push-pull flow scheduling, Figure 5 E and J show relatively flat profiles despite the GBM model still containing discernable diminished penetrability of the GBM. Figure 5 A and F show the velocity profile of the 300-600 minute flow scheme illustrating the GBM flow blockage. Induced flow does not penetrate below the tumor indicating that a variation in z-height of the catheters may be necessary to effect push-pull of the drug across all components of the GBM.

## Discussion

Utilizing current neuro-reconstruction/analysis software’s combined with a differential equation’s solver, we designed and tested a pipeline for simulating of an individual’s hemisphere from their unique MRI. In this model for healthy and diseased brains, we defined a variety of parameters (Table 1) in the simulation software to create a platform that simulates key characteristics known to impact molecular diffusion, including porosity and permeability. With this construction, we were able to simulate convection enhanced delivery of drugs, biomarker uptake, and multi-catheter optimization of drug delivery.

Our simulation builds on previous efforts of simulating interstitial fluid flow and solute dispersal. Notable in previous simulations, minimal pia mater spatial replication was attempted. To model various fluid permeation diffusion properties, the simplest brain topography models have typically used 2D slices or thin 3D slices. Vidotto et al. used thin 3D slices to simulate the microstructure of brain matter, while Shahim et al. used 2D slices to simulate the anisotropy to brain matter [30], [31]. Other popular mechanisms of brain simulations include those proposed by ‘Therataxis, Baltimore, MD, USA and BrainLAB, Feldkirchen’ and have been used in prior studies to simulate drug delivery to the brain [32]–[34]. Brainlab’s simulation focuses on utilizing patient-observed solute diffusivity in brains developing fitting equations. These equations incorporate factors such as cellular/vascular drug uptake, diffusivity correcting factors, and bulk flow. These traits were developed into a governing equation by observing patient data. The more common simulation scheme, however, is a geometric construction with smooth surfaces that roughly approximates the shape of the brain beneath the skull [9], [35]–[38]. While these models have laid the foundation for brain simulations, they are an imperfect representation of each patient’s unique cerebral topography, thus decreasing the translational applicability of such models. A promising diffusion study by Zhan et al. simulated a more detailed topographical brain [39] by distinguishing the brain’s gyri/sulci in the pial; however, it did not divide the left and right hemispheres of the brain and the sulci were not well defined. Without the impedances of the central wall, diffusion would be considerably uniform relative to the many perturbations present *in vivo*.

Establishing the parenchymal space for simulations is an essential step for evaluating drug dispersal; however, multiple critical traits impacting have not been included. These factors include membranes, drug and biomarker activity, and additional structures. Membranous features such as the pial surface, ventricular to parenchymal interface, and tractography interfaces regulate solute and momentum flow and induce capillary perturbations. Meanwhile drug and biomarker concentration levels are subject to cellular uptake and degradation rates, vascular uptake, and local variations of production for biomarkers within GBMs [40]. Bulk interstitial CSF flow is driven by a ventricular circuit perfusing the brain tissue requiring patient specific ventricular construction for replicability for simulations. While these traits limit the current work in its applicability for accurately representing drug dispersal in the brain, our future work will be to remedy these limitations.

## Conclusion

We have demonstrated that a patient’s MRI can be converted into a volumetric scan and precisely edited to generate an unprecedented *in silico* brain mesh model with detailed and individualized 3D topography. for diffusion computational analyses. This model has the potential to be deployed as a platform for surgical planning, drug and biomarker diffusions studies, and evaluations of implantable neural devices and techniques. Prior to these applications, future studies will need to parameterize more physiologically relevant traits, including more detailed and complex anatomy, interstitial fluid flow, material properties, anisotropic material properties, and generation and consumption rates of tumor biomarkers and drugs.

## Acknowledgement

MRK and TB conceived and supervised the project. MRK acknowledges the funding support of the VPRI startup fund. LS provided and SO developed and guided the computational support for the conversion of MRI to digital usable format for further analysis. LR conducted the conversion procedures. The MRI data was acquired from NeuroImaging Tools & Resources Collaboratory (NITRC) Dallas Lifetime Study subjects, funded by NIH grant numbers 2R44NS074540 and 1U24EB023398. DO aided the analysis. CRC contributed to the writing and analysis. CS designed, executed, and experimented the project; analyzed the results; and wrote the Manuscript.

## Supplemental Information

**Table S1.** Computational Meshing Results.

| CED No Tumor |  | CED Tumor |  | Four Probe No Tumor |  | Four Probe Tumor |  |
| --- | --- | --- | --- | --- | --- | --- | --- |
| Tetrahedron | 1896373 | Tetrahedron | 1889469 | Tetrahedron | 3963903 | Tetrahedron | 2620744 |
| Pyramid | 5839 | Pyramid | 5839 |  |  |  |  |
| Prism | 182797 | Prism | 182797 |  |  |  |  |
| Triangle | 110945 | Triangle | 112361 | Triangle | 184176 | Triangle | 148156 |
| Quad | 453 | Quad | 453 |  |  |  |  |
| Edge element | 5479 | Edge element | 5601 | Edge element | 9271 | Edge element | 6193 |
| Vertex element | 332 | Vertex element | 338 | Vertex element | 398 | Vertex element | 430 |
| Total | 2202218 | Total | 2196858 | Total | 4157748 | Total | 2775523 |

**Figure S1.**
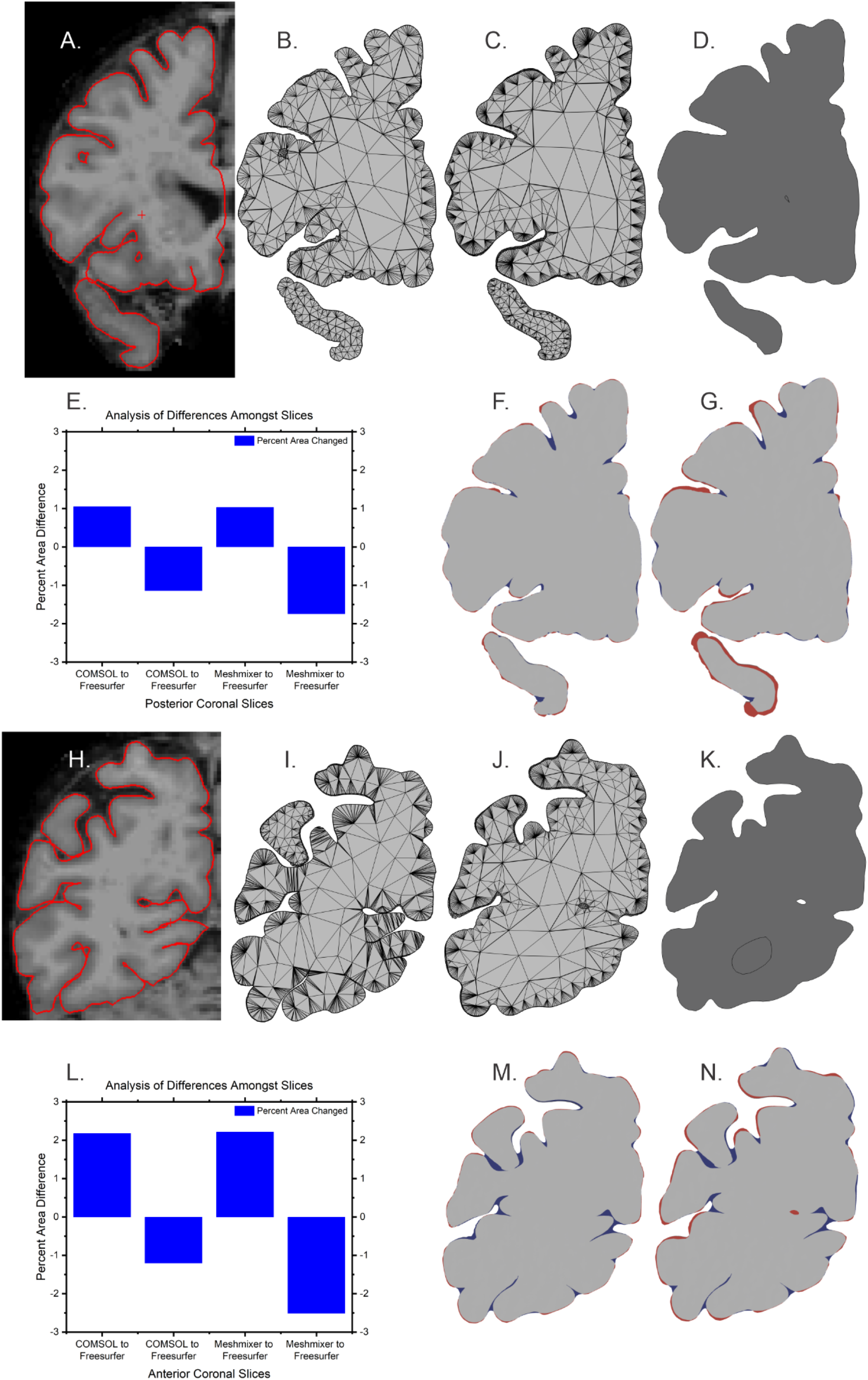
Comparison of coronal slices to demonstrate conformity of replication during processing. A. and H. reconstructed MRI slices with pial surface highlighted in red for anterior and posterior regions. B. and I FreeSurfer generated volumetric data of pial surface layer. C. and J Meshmixer’s computational mesh reconstruction of pial surface layer. D. and K. COMSOL Multiphysics computational mesh of pial surface layer after simulation. E. and L. Plot of area added and subtracted to the original FreeSurfer slice. F. and M. Slice showing area added and subtracted to original FreeSurfer mesh by the Meshmixer edited mesh. The blue regions show areas added and the red show areas subtracted from the FreeSurfer slice. G. and N. Slice showing the blue added and the red subtracted regions of the FreeSurfer slice by the COMSOL slice.

